# Volatile metabolites in lavage fluid are correlated to Valley fever disease severity in murine model lung infections

**DOI:** 10.1101/2022.10.02.510537

**Authors:** Emily A. Higgins Keppler, Marley C. Caballero Van Dyke, Heather L. Mead, Douglas F. Lake, D. Mitchell Magee, Bridget M. Barker, Heather D. Bean

**Affiliations:** School of Life Sciences, Arizona State University, Tempe, Arizona, USA; Center for Fundamental and Applied Microbiomics, The Biodesign Institute, Arizona State University, Tempe, Arizona, USA; Pathogen and Microbiome Institute, Northern Arizona University, Flagstaff, Arizona, USA; Center for Personalized Diagnostics, The Biodesign Institute, Arizona State University, Tempe, Arizona, USA

## Abstract

*Coccidioides immitis* and *Coccidioides posadasii* are soil-dwelling fungi of arid regions in North and South America that are responsible for Valley fever (coccidioidomycosis). Forty percent of patients with Valley fever exhibit symptoms ranging from mild, self-limiting respiratory infections, to severe, life-threatening pneumonia that requires treatment. Misdiagnosis as bacterial pneumonia commonly occurs in symptomatic Valley fever cases, resulting in inappropriate treatment with antibiotics, increased medical costs, and delay in diagnosis. In this study, we explored the feasibility of developing breath-based diagnostics for Valley fever using a murine lung infection model. To investigate potential volatile biomarkers of Valley fever that arise from host-pathogen interactions, we infected C57BL/6J mice with *C. immitis* RS and *C. posadasii* Silveira via intranasal inoculation. We measured fungal dissemination and collected bronchoalveolar lavage fluid (BALF) for cytokine profiling and for untargeted volatile metabolomics via solid phase microextraction (SPME) and two-dimensional gas chromatography coupled to time-of-flight mass spectrometry (GC×GC-TOFMS). We identified 36 volatile organic compounds (VOCs) that were significantly correlated to cytokine abundances and clustered mice by disease severity. These 36 VOCs were also able to separate mice with a moderate to high disease severity by infection strain. The data presented here show that *Coccidioides* and/or the host produce volatile metabolites that may yield biomarkers for a Valley fever breath test that can detect Coccidioidal infection and provide clinically relevant information on disease severity.

**IMPORTANCE:** Coccidioidomycosis, or Valley fever, is a fungal disease endemic to the North and South American arid regions. Forty percent of individuals infected with Valley fever will exhibit symptoms consistent with community-acquired pneumonia. However, misdiagnosis frequently occurs in these cases, resulting in inappropriate treatment with antibiotics, increased medical costs, and delay in receiving an accurate diagnosis. Herein, we used a murine lung infection model as a step towards developing a breath-based diagnostic for Valley fever. We infected C57BL/6J mice with *C. immitis* RS and *C. posadasii* Silveira and collected bronchoalveolar lavage fluid for untargeted volatile metabolomics. We observed that volatile metabolites in the bronchoalveolar lavage fluid of Cocci-inoculated mice were significantly correlated to disease severity, as measured by immune response. The data presented here show that *Coccidioides* and/or the host produce volatile metabolites that may yield biomarkers for a Valley fever breath test.

## INTRODUCTION

*Coccidioides immitis* and *C. posadasii* are soil-dwelling fungi of the arid regions of North and South America that cause the fungal infection coccidioidomycosis, or Valley fever, when arthroconidia are inhaled. One-half to two-thirds of individuals infected with *Coccidioides* remain asymptomatic or exhibit mild symptoms and they resolve their clinical disease without seeking medical attention, retaining long-lived immunity (1). Symptoms develop in approximately forty percent of Valley fever cases ranging from mild, self-limiting respiratory infections, to severe, life-threatening pneumonia, and a small percentage will result in disseminated disease (2). The most common diagnosis is acute or subacute pneumonic illness, which usually does not require treatment; instead, patients are monitored until the infection is resolved on its own (1). Conversely, patients with extensive spread of infection or who are at high risk of complications due to immunosuppression or other preexisting conditions require a variety of treatment strategies including antifungal therapy, surgical debridement, or both (1).

Misdiagnosis occurs frequently in symptomatic Valley fever. Without testing, it is often mistaken for a bacterial pneumonia, and inappropriately treated with antibiotics. The only conclusive diagnosis of Valley fever requires a positive fungal culture or histopathology showing spherules or endospores within lung tissue (3). However, due to the invasive nature of obtaining lung tissue, most patients are diagnosed through serological testing, but low sensitivities contribute to diagnostic delays. Enzyme immunoassay (EIA), immunodiffusion (ID), and complement fixation (CF) tests are all commercially available; however, EIA tests for IgG and IgM range in sensitivity from 47-87% and 22-61%, respectively, and ID and CF tests are between 60-91% and 65-98% sensitive, respectively (2). The combination of insufficient testing for fungal pneumonias and insufficient sensitivities in the existing diagnostics lead to more than a one month delay in diagnosis for 46% of Valley fever patients in the endemic region of Phoenix, Arizona, United States, significantly increasing medical costs (4). Surveillance in non-endemic states noted that 70% of Valley fever patients were diagnosed with another condition before being tested for coccidioidomycosis, with a median delay in diagnosis of 38 days (5).

Breath-based diagnostics are increasingly being pursued as a means for diagnosing respiratory infections. Animal model respiratory infections have shown that the combination of pathogen and host volatile metabolites have high diagnostic accuracy for detecting and identifying lung infection etiology (6-15). Further, clinical studies of community-acquired pneumonia, ventilator-associated pneumonia, and chronic lung infections have shown that volatile biomarkers can sensitively detect and identify the bacterial and fungal etiologies through the analysis of sputum, bronchoalveolar lavage fluid (BALF), and breath volatiles (16-19). In this study we explored the feasibility of developing breath-based diagnostics for Valley fever using a murine lung infection model. To demonstrate that *Coccidioides* volatile biomarkers can be detected *in vivo*, and to identify volatile biomarkers of Valley fever that arise from host-pathogen interactions, we infected mice with *C. immitis* and *C. posadasii* and collected BALF for untargeted volatile metabolomics via two-dimensional gas chromatography coupled to time-of-flight mass spectrometry (GC×GC-TOFMS). Since an antibody response is often delayed, we hypothesized that *Coccidioides* and the host would produce volatile metabolites in response to infection that can be utilized as sensitive and specific biomarkers for Valley fever infections.

## RESULTS

### *Coccidioides* infected C57BL/6J mice exhibit a gradient of disease severity

Six mice from each infection group, *Coccidioides immitis* strain RS and *C. posadasii* strain Silveira (Sil), and four control mice that were sham-infected with phosphate-buffered saline (PBS) were sacrificed 10 days post-inoculation. Bronchoalveolar lavage fluid (BALF) was collected and analyzed for the abundances of 26 immune markers, and disseminated disease was assessed by quantifying the number of fungal colonies cultured from the mouse spleen and brain. All four PBS-inoculated mice produced very low levels of cytokines, but the Cocci-inoculated mice produced a wide range of immune responses to the fungus (Figure 1; Supplementary Table S1). Two of the Cocci**-**inoculated mice, RS 1 and Sil 6, produced a high abundance of cytokines at 39- and 38-fold greater than PBS mice, respectively. Two mice, RS 5 and 6, produced very low levels of cytokines, equivalent to the abundances detected in the BALF of PBS mice, though the cytokine profiles of the infected and uninfected mice were different (Supplementary Figure S1). The remaining Cocci-inoculated mice produced a gradient of cytokine concentrations from 3-to 23-fold greater than controls. Generally, fungal dissemination correlated with the degree of cytokine response, independent of the cytokine profile. Fungus was recovered from the spleen and brain in the three RS mice with the highest cytokine responses and in the spleen of all six of the Silveira mice, four of which also had detectable dissemination to the brain (Figure 1, Supplementary Table S1). Performing BAL on the mice precluded direct measurements of fungal lung disease via histology or tissue homogenization and plating; therefore, we utilized the combination of cytokine concentrations and fungal disseminations as proxies for coccidioidomycosis disease severity in this study. The correlation between fungal dose and immune response in murine model lung infections of coccidioidomycosis has been previously reported (20, 21).

**Figure 1.**
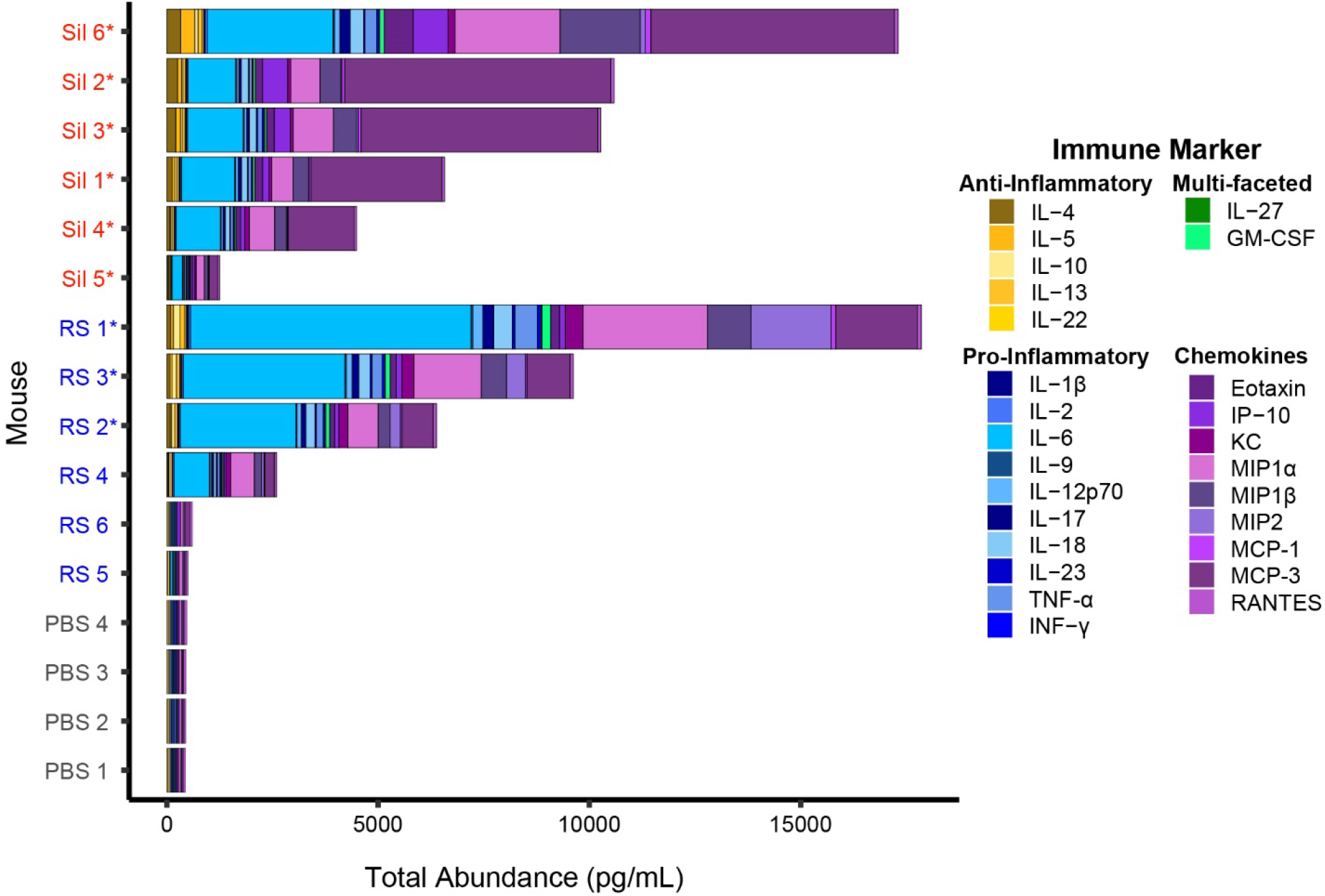
Abundances of 26 cytokines in individual mice inoculated with *C. immitis* strain RS (blue) and *C. posadasii* strain Silveira (Sil; red), or sham-inoculated with PBS (gray). Mice with disseminated disease are indicated with an asterisk (*). Immune markers are color-coded by type.

Similar to other studies characterizing the murine immune responses to RS and Silveira (22-25), we detected differences in the cytokine profiles of mice infected with these two strains; e.g., we found that IL-6 is more abundant in RS-infected mice and MCP-3 is more abundant in Silveira (Figure 1). However, the dominating pattern that we observe in these data is based on the differences in disease severity. Principal component analysis (PCA), using the 12 Cocci-inoculated and 4 PBS mice as observations and 26 cytokines as variables, shows that the mice separate by disease severity on PC 1 (representing 88.4% of the total variance), with the two mice with the lowest severity, RS 5 and RS 6, clustering with the PBS controls (Figure 2). The species-level differences in the cytokine profiles are represented by PC 2, which captures only 6.9% of the total variance. A hierarchical clustering analysis (HCA) of the mice based on their relative cytokine abundances generates two clusters, with one cluster containing the mice with moderate to high disease severity and the other containing the PBS mice and the three Cocci-inoculated mice with the lowest severity (Supplementary Figure S2). Within the moderate-to-high disease severity cluster there is not a clear separation in the mice by fungal strain; rather, the clustering is based upon overall cytokine abundances.

**Figure 2.**
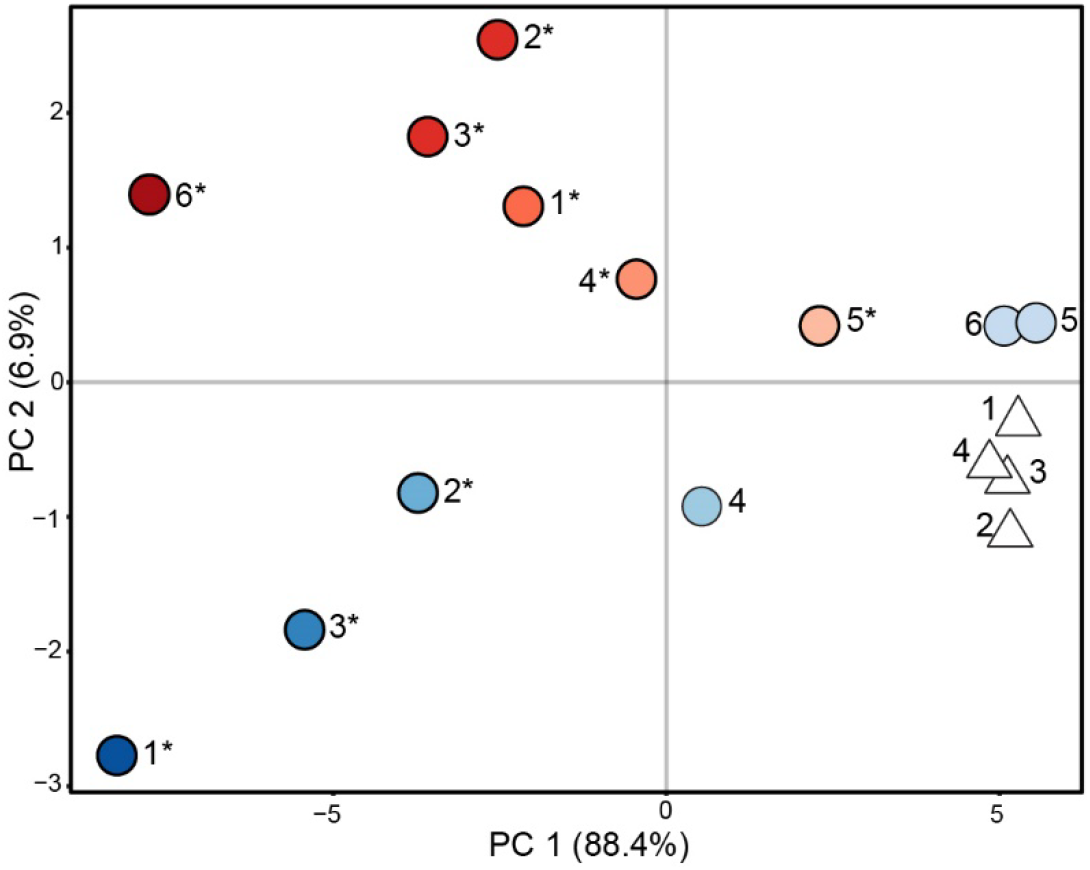
Principal component analysis (PCA) score plot using the abundances of 26 cytokines as features, and mice inoculated with *C. immitis* RS (blue circles), *C. posadasii* Silveira (red circles) or PBS (white triangles) as observations. The color gradient, darkest to lightest, indicates total cytokine abundance, highest to lowest; disseminated disease is indicated with an asterisk (*).

### Murine coccidioidomycosis volatilome and its correlation to disease severity

The volatile compounds present in the BALF samples were collected using solid phase microextraction (SPME) and analyzed using two-dimensional gas chromatography coupled to time-of-flight mass spectrometry (GC×GC-TOFMS). After contaminant removal and data clean-up we detected 91 VOCs (Supplemental Table S2), of which 26 were identified at a level 1 or 2 (26) and were assigned putative compound names based on mass spectral and chromatographic data (Table 1). Of the 26 named compounds, 19 have been previously associated with human and environmental fungal pathogens, including decanal, which was detected in the *in vitro* cultures of *Coccidioides* spp. (27) (Table 1). For the unnamed compounds, 16 were identified at level 3 and assigned chemical classification based on a combination of mass spectral and chromatographic data (Supplemental Table S2).

**Table 1.**
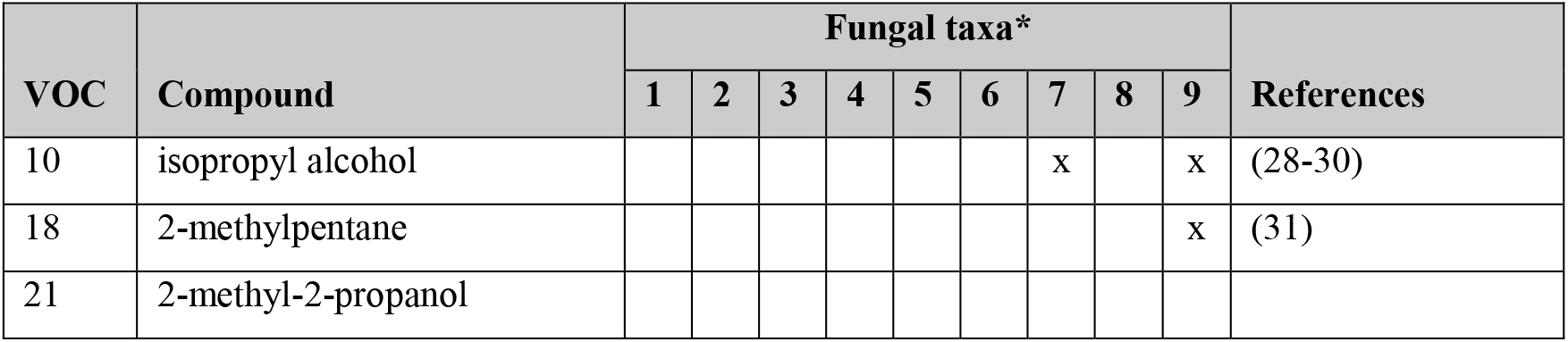
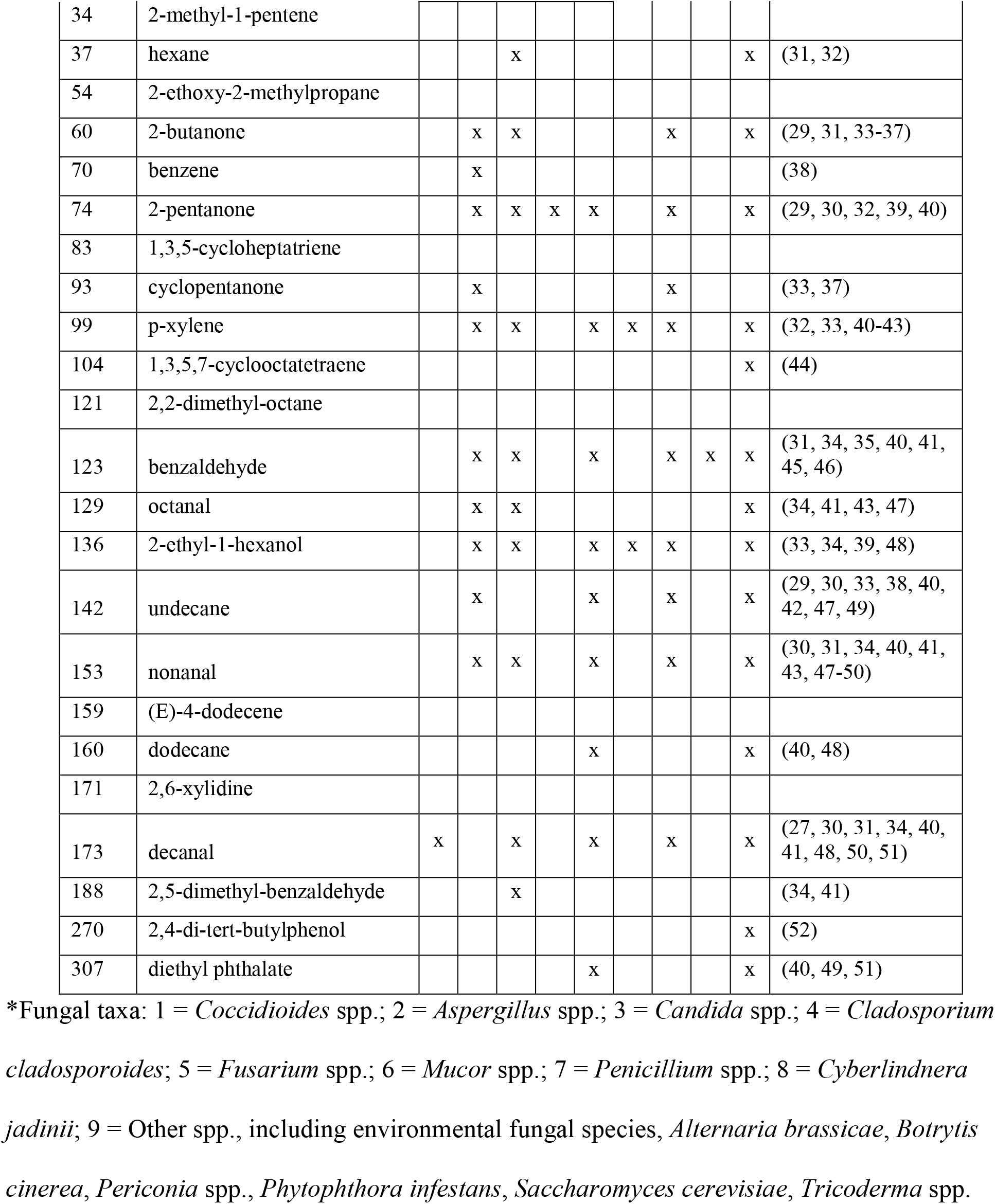
Named volatiles detected in the BALF of *Coccidioides*-infected mice and their previous reports in other fungal taxa.

To determine the relationship between *Coccidioides* infection and the BALF volatilome, we performed PCA using the 91 VOCs as features and the mice as observations and detected a trend toward the mice separating on PC 1 based on disease severity (Supplementary Figure S3). Using all of the VOCs, however, only 18.3% of the variance is represented by PC 1 and the separation by disease severity was weaker than we observed for the cytokines, likely because only a subset of the BALF VOCs are altered by *Coccidioides* infection. A Kendall correlation between the 26 immune markers and 91 volatiles showed that 36 VOCs (40% of the total volatilome) were significantly correlated to at least one cytokine (–0.3 > τ > 0.3; *P* < 0.05; Supplementary Figure S4; Figure 3). As seen in a previous study on the relationship between infection VOCs and immune response in murine model lung infections (10), each VOC is either positively (or negatively) correlated to the full suite of cytokines, versus the VOCs having mixed patterns of positive and negative correlation. Therefore, the abundances of the immune-correlated VOCs are related to overall disease severity rather than to specific immune pathways.

**Figure 3.**
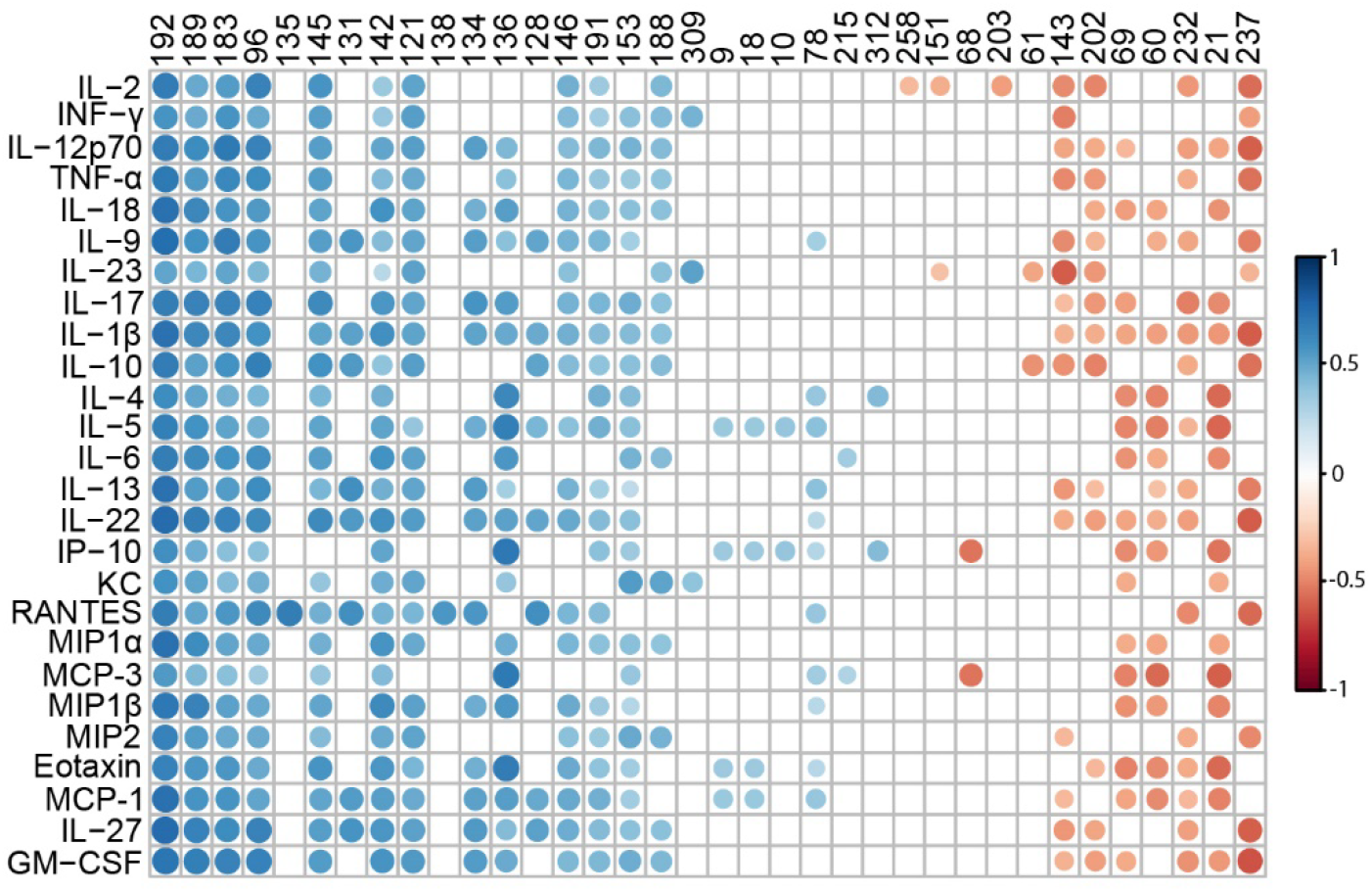
Kendall correlation plot of the 36 volatile organic compounds (VOCs) (columns), that are significantly correlated with at least one of the 26 cytokines (rows), produced by the 12 Cocci-inoculated and 4 PBS-inoculated control mice. Volatiles are ordered by mean correlation from most positive (left) to most negative (right). Circles indicate statistically significant correlations (*P* < 0.05), with positive correlations in blue (τ > 0.3), negative correlations in red (τ< –0.3), and darker colors and larger sizes indicating a stronger correlation.

When we perform PCA using only the 36 immune-correlated VOCs as variables, we observe that twice as much variance is explained by disease severity on PC 1 compared to the PCA with all 91 VOCs (Figure 4a, Supplementary Figure S3). The loading pattern of the VOCs on PC 1 (Figure 4b) mirrors the Kendall correlations, with the 12 VOCs that are negatively correlated to cytokines driving the separation of the controls and the low disease severity mice onto PC 1 > 0 and the remaining 24 positively correlated VOCs loading on PC 1 < 0 where the moderately-to-highly infected mice cluster. On PC 2, four volatiles (21, 60, 68, and 69) separate the control mice from the five Cocci-inoculated mice with the lowest disease severity (Figure 4b). Additionally, mice with moderate to high disease severity separate on PC 2 based on the strain with which they were infected.

**Figure 4.**
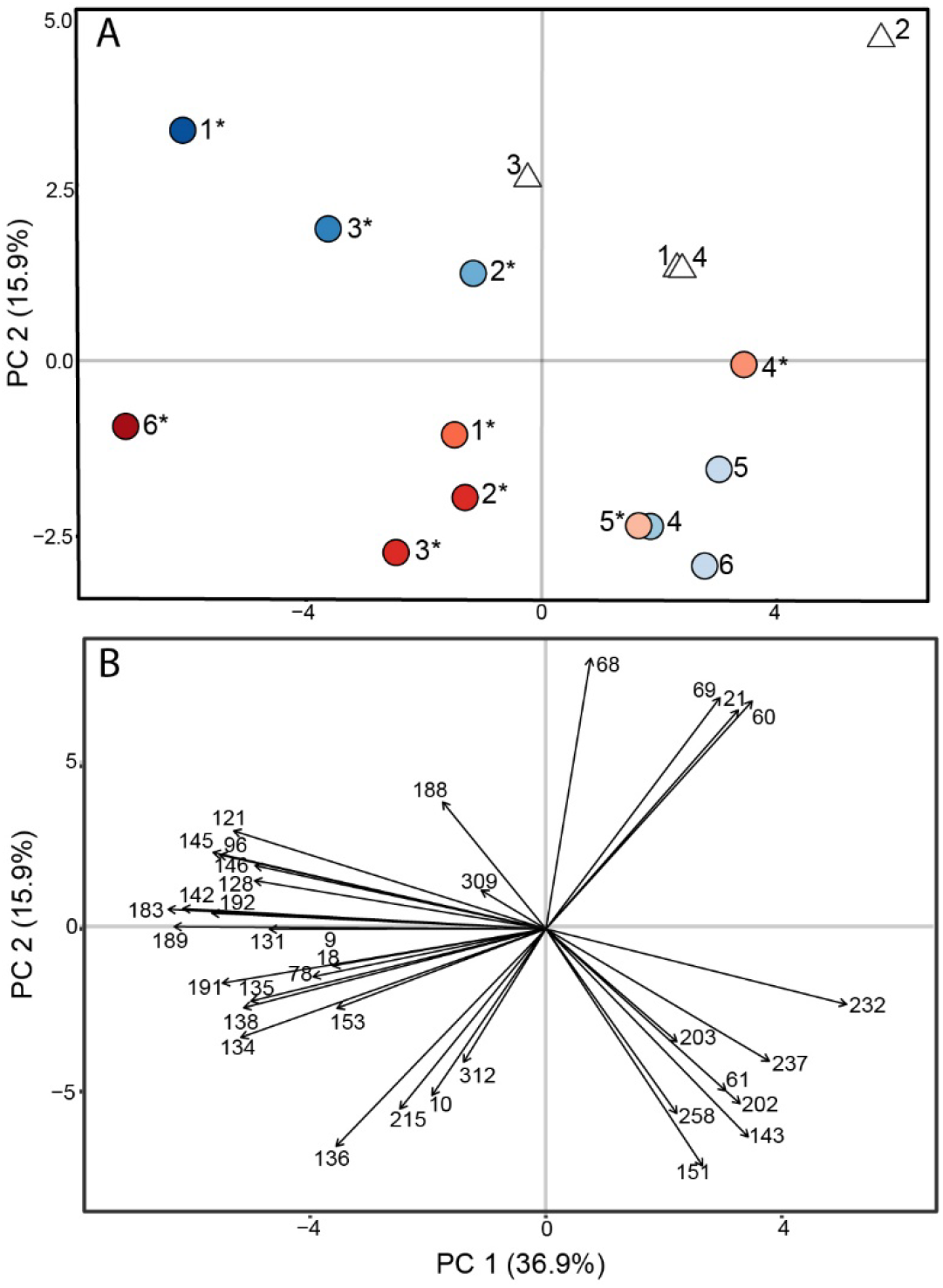
Principal component analysis score plot (**A**) and loading plot (**B**) using 36 immune-correlated volatile organic compounds (VOCs) as features, and mice inoculated with *C. immitis* RS (blue circles), *C. posadasii* Silveira (red circles) or PBS (white triangles) as observations. The color gradient, darker to lighter, indicates total cytokine abundance, higher to lower; disseminated disease is indicated with an asterisk (*).

HCA using the 36 immune-correlated VOCs as variables recapitulates key features of the PCA with mice separating into two main clusters based on disease severity, and the mice with moderate to high disease severity clustering by strain of infection (Figure 5). Additionally, the 36 volatiles divided into two clusters: volatiles that are positively correlated (τ > 0.3) to cytokine production in one cluster, and the negatively correlated (τ < –0.3) volatiles in another. This division is also what drives the variance across PC 1 (Figure 4b). Six of the 36 immune-correlated VOCs are uniquely produced by mice with moderate to high disease severity (VOCs 9, 18, 121, 146, 192, and 312; Figure 5 and Supplemental Table S2), with three of them only detectable in the BALF of Sil 6 and RS 1, who had the most severe disease.

**Figure 5.**
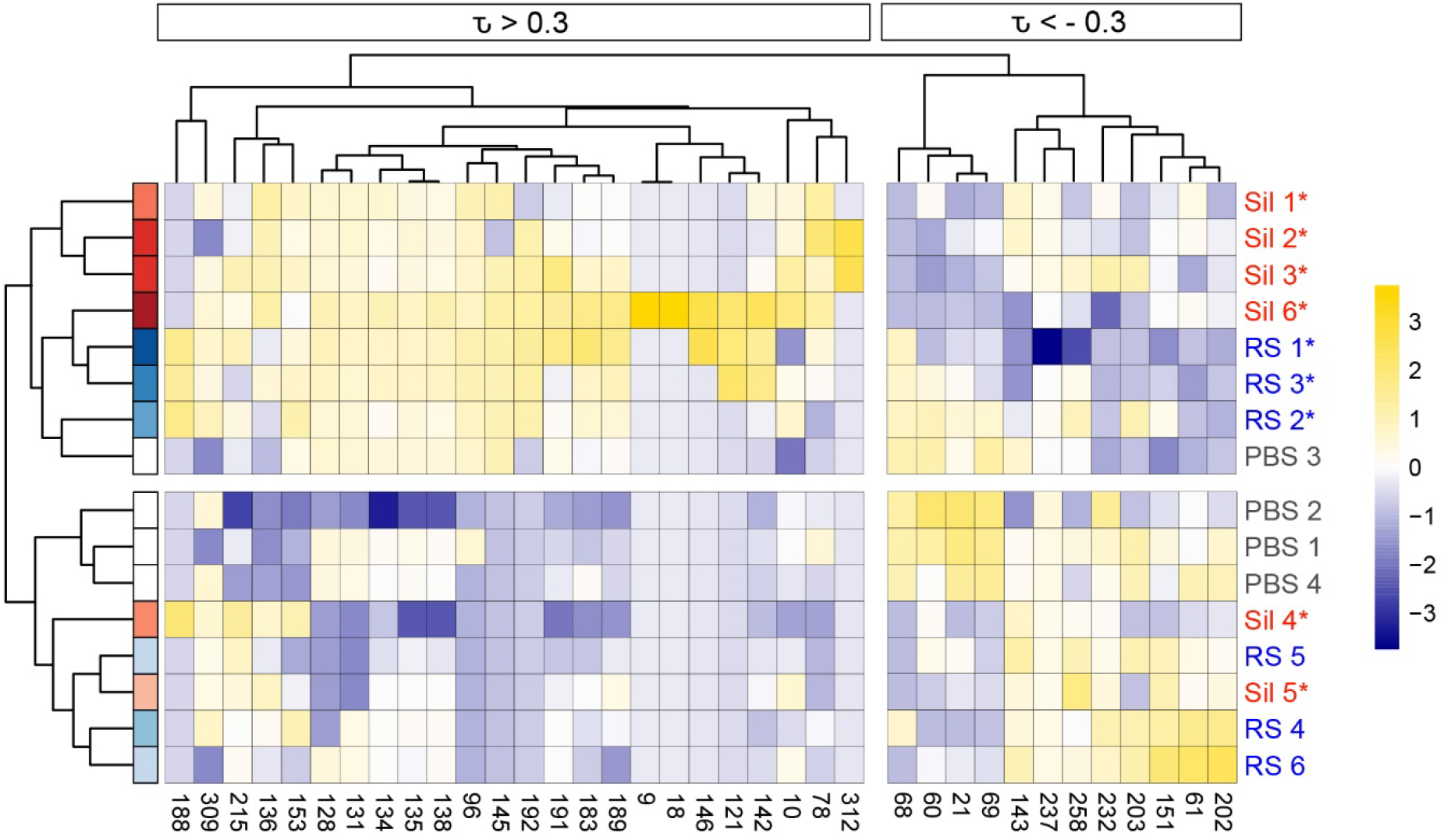
Hierarchical clustering analysis (HCA) of 12 Cocci-inoculated and 4 PBS mice (rows) based on the relative abundance of 36 immune-correlated volatile organic compounds (VOCs) (columns). Clustering of mice and VOCs uses Pearson correlations with average linkage. Mice are color-coded by species (blue = *C. immitis* RS; red = *C. posadasii* Sil) and a color gradient indicating total cytokine abundance, with darker color meaning higher abundance; disseminated disease is indicated with an asterisk (*). Kendall correlation between volatiles and cytokines is noted above the dendrogram, with τ > 0.3 for positive correlations and τ < -0.3 for negative. Volatiles abundances (mean-centered and scaled to unit variance) are represented in the heat map.

## DISCUSSION

For this study, we sought to characterize the volatilome of a murine model of coccidioidomycosis lung infection. We examined the BALF of mice infected with *C. immitis* strain RS and *C. posadasii* strain Silveira as well as sham-infected PBS mice. We initially anticipated that all Cocci-inoculated mice would have comparable cytokine levels in response to infection; however, what we found was a gradient of disease severity across the mice, represented by cytokine abundance and fungal dissemination (Figure 1). While we strove to give each mouse the same dose of arthroconidia, due to the intranasal route of administration, it is possible that some mice swallowed a portion of the inoculum, reducing the overall dose received. However, we can see that even the mice with the lowest levels of BALF cytokines, RS 5 and 6, have a different cytokine profile than that of the PBS mice, indicating that there was a weak immune response to fungal exposure (Figure S1). While C57BL/6J mice are inbred, they are not genetically identical, and other mouse model studies have observed differences between individual mice given the same intranasal dose of a pathogen. Zhu et al. quantified the bacterial cell counts of homogenized lung tissue at multiple time points after 72 C57Bl/6J mice were infected with either *Pseudomonas aeruginosa* or *Staphylococcus aureus* (8). Six mice were sacrificed at each time point, and while the mice had very similar bacterial cell counts for the first four time points post inoculation (i.e., CFU/lung within one order of magnitude, up to 48 h p.i.), by 72 h and 120 h p.i. differences in the response to infection emerged; some mice had cleared the infection (i.e., no viable bacteria recovered), some had bacterial cell counts in their lungs ranging from 10^3^ - 10^8^, and a few mice succumbed to their infections. Franchina et al. observed a 100-fold difference in bacterial cell counts in the BALF of C57BL/6J mice infected with *Mycobacterium tuberculosis* at either 24 or 48 h post infection (13). Additionally, they observed a 100,000-fold difference in the number of neutrophils and macrophages present in the BALF of mice at each time point. Therefore, the variability in disseminated disease and in BALF cytokines that we observed within the RS- and Sil-inoculated cohorts are consistent with other murine pneumonia model studies in this mouse strain.

Of the 26 named compounds that we detected in the BALF of the infected mice, only one, decanal, was previously detected and identified in the *in vitro* cultures of *Coccidioides* spp. (27). The lack of *in vitro* volatiles in this present study is not surprising, given the changes that *Coccidioides* undergoes within a mammalian host (53), and the significant contributions that the host makes to the infection volatilome (10). A prior mouse model study and two human studies on ventilator-associated pneumonia reported that only about one quarter to one third of bacterial *in vitro* volatiles are also detected in the breath of the host (7, 41, 54). Thus, while this study provides the proof-of-concept that the lung volatiles of *Coccidioides* infected hosts significantly differ from those without infection, the most direct route to the development of breath biomarkers for Valley fever will be in human clinical studies.

Forty percent of the volatiles that we detected in BALF are correlated to disease severity, with the volatiles that are negatively correlated to cytokine concentrations leading to separation of the low severity and control mice, while the positively correlated volatiles drive the separation of the more severely diseased mice by the strain of infection (Figures 4 and 5). Because this study only included one strain from each species, we are unable to extrapolate the strain-specific separation of the infection volatiles that we observed here to the likelihood that there are inherent species-specific differences between volatiles from Valley fever caused by *C. immitis* vs. *C. posadasii* infection. However, our prior analysis of the *in vitro* volatilome of six strains each of *C. immitis* and *C. posadasii*, which included strains RS and Silveira, suggest that it is unlikely that we would observe inter-species differences in infection volatiles because intra-species volatilomes are highly diverse (27). Additionally, previous *Coccidioides* mouse studies have shown a variation in immune response between mice and *Coccidioides* strains, suggesting that the host’s contribution to Valley fever volatile profile may also vary by strain (55, 56). While both RS and Silveira are the type strains for their respective species, they both have characteristics that make us cautious to generalize these results to other strains of *C. immitis* or *C. posadasii*. Recent whole genome sequencing of RS revealed that it is a hybrid strain with approximately 20% of its genome from *C. posadasii* (57). Further, our *in vitro* volatilome study suggests RS has a metabolism that differs from many other *Coccidioides* strains, especially in the mycelial form where it failed to produce many key volatile compounds that differentiate that lifeform from spherules (27). Silveira is often selected in vaccine challenges due to its high virulence in mice, which may not be representative of all *C. posadasii*. To understand the variability of Valley fever infection volatiles and the impact that may have on the development of a Valley fever breath test, additional *Coccidioides* strains need to be included in future murine model infection studies.

While very small differences in cytokine abundances separated the PBS control mice from RS 5 and RS 6, which had very low levels of disease (Figure 2), a higher minimum threshold of disease severity was required before the volatiles clustered infected mice away from the controls (Figures 4 and 5). This may be due to the limits of detection for VOCs from 200 μL of BALF, and by collecting higher BALF volumes or sampling and concentrating multiple breaths, we would be able to detect lower levels of Valley fever disease using VOCs. Clinically it is recommended that patients with primary pulmonary coccidioidomycosis are initially monitored rather than treated with antifungals, as most patients resolve their infection without intervention (3). It is further advised that treatment with antifungals be reserved for patients with underlying immunosuppression, substantial comorbidities, or particularly severe disease (2).

Therefore, the ideal Valley fever diagnostic will be able to distinguish not only presence or absence of *Coccidioides* but also level of infection, thereby limiting unnecessary antifungal treatment. The data presented here suggest it will be possible to develop a Valley fever breath test that will be sensitive enough to indicate the degree of disease severity. With such a diagnostic, clinicians can make an informed decision on treatment, reserving antifungals for the patients who will not be able to clear the infection on their own, while not over treating individuals who will have self-limiting disease. It will be important going forward to not only define the breath biomarker threshold for initiating treatment, but also to recognize this prescribing threshold will require a precision medicine approach; a lower disease severity in patients who have co-morbidities that increase their risk of prolonged infection (e.g. TNF-inhibitors, chemotherapy, solid organ transplant) may necessitate the use of antifungal medications.

## CONCLUSIONS

C57BL/6J mice inoculated with *C. immitis* RS or *C. posadasii* Silveira produced a gradient of disease severity and cytokine abundance. We identified 36 VOCs that were significantly correlated to cytokine production, and these specific volatiles clustered mice by disease severity. These 36 VOCs were also able to separate mice with a moderate to high disease severity by infection strain. The data presented here show that *Coccidioides* and/or the host produce volatile metabolites that may yield biomarkers for a Valley fever breath test that can detect the fungus and provide clinically-relevant information on disease severity.

## METHODS

### Mice

Female C57BL/6J mice (The Jackson Laboratory, Bar Harbor, ME) 6-8 weeks of age were used for these studies. Mice were housed according to NIH guidelines for housing and care in a biosafety level 3 animal laboratory. All procedures were approved by the Institutional Animal Care and Use Committee (protocol number 16-011) of Northern Arizona University.

### Pulmonary Coccidioidal Infections

The *Coccidioides* isolates used in this study were the type strains *C. immitis* strain RS (ATCC^®^ catalog no. NR-48942; NCBI accession no. AAEC00000000.3) and *C. posadasii* strain Silveira (ATCC^®^ catalog no. NR-48944; NCBI accession no. ABAI00000000.2). Mice were anesthetized with ketamine/xylene (80/8 mg/kg) and intranasally inoculated with 100 arthroconidia of *C. immitis* strain RS (*n*=6) or *C. posadasii* strain Silveira (*n*=6) suspended in 30 μL phosphate-buffered saline (PBS), as described previously (22, 58). Control mice were inoculated with PBS alone (*n*=4). The mice were sacrificed at day 10 post-inoculation. The lungs were rinsed with 2 mL of PBS to collect bronchoalveolar lavage fluid (BALF), which were filtered with 0.22 μm Ultrafree^®^ -MC centrifugal filter devices with Durapore^®^ membrane (MilliporeSigma, Burlington, MA). One milliliter of each BALF sample was stored at –80°C for volatilomics analysis. Halt™ Protease Inhibitor Cocktail (10 μL/mL) was added to the remainder of each BALF sample for cytokine analysis. Spleen and brain were homogenized in 1 ml of sterile PBS followed by culture of 10-fold dilutions of each tissue on 2X GYE agar (2% glucose (VWR™, USA), 1% yeast extract (BD™, Franklin Lakes, New Jersey, USA, and 1.5% bacteriological agar (Difco, USA)) to assess fungal dissemination.

### Cytokine Analysis

Cytokine levels in the BALF samples were examined using the cytokine mouse magnetic Invitrogen™ 26-Plex ProcartaPlex™ Panel 1 (25 μL of sample; ThermoFisher Scientific) per manufacturer’s instructions using a 2 h incubation. Samples were read using a MAGPIX^®^ multiplex system (Luminex^®^ Corporation, Austin, TX) with Luminex^®^ xPONENT^®^ 3.0 software.

### Volatile metabolomics analysis by SPME-GC×GC-TOFMS

The BALF samples were allowed to thaw at 4°C overnight, and then split into technical triplicates of 200 μL that were transferred and sealed into sterilized 2 mL GC headspace vials with Supelco^®^ PTFE/silicone septum magnetic screw caps (Sigma-Aldrich^®^, St. Louis, MO). Samples were randomized for analysis. Volatile metabolites sampling was performed by solid phase microextraction (SPME) using a Gerstel^®^ MPS Robotic Pro MultiPurpose autosampler directed by Maestro^®^ software (Gerstel^®^, Inc., Linthicum, MD). Sample extraction and injection parameters are provided in Table S3 (see Autosampler Method). Volatile metabolite analysis was performed by two-dimensional gas chromatography−time-of-flight mass spectrometry (GC×GC– TOFMS) using a LECO^®^ Pegasus^®^ 4D and Agilent^®^ 7890B GC (LECO^®^ Corp., St. Joseph, MI). Chromatographic, mass spectrometric, and peak detection parameters are provided in Table S3 (see GC×GC Method and Mass Spectrometry Method). An external alkane standards mixture (C_8_ – C_20_; Sigma-Aldrich^®^) was sampled multiple times for calculating retention indices (RI). The injection, chromatographic, and mass spectrometric methods for analyzing the alkane standards were the same as for the samples.

### Processing and analysis of chromatographic data

Data collection, processing, and alignment were performed using ChromaTOF software version 4.71 with the Statistical Compare package (LECO^®^ Corp.), using the parameters listed in Table S3 (see Data Processing Method). Peaks were assigned a putative identification based on mass spectral similarity and retention index (RI) data, and the confidence of those identifications are indicated by assigning levels 1 to 4 (1 highest) (26). Peaks at level 1 were identified based on mass spectral and RI matches with external standards. Peaks at level 2 were identified based on ≥ 800 mass spectral match by a forward and reverse search of the NIST 14 library and RIs that are consistent (< 7% error) with a calculated 624 RI using the following equation: mean NIST nonpolar RI x 1.0517 + 14.468; this equation was obtained based on the linear relationship of RIs observed between a 624Sil column and nonpolar stationary phase (59). Level 1, 2, and 3 compounds were assigned to chemical functional groups based upon characteristic mass spectral fragmentation patterns and second dimension retention times, as previously described (60). Level 4 compounds have mass spectral matches < 600 or RI that do not match previously published values and are reported as unknowns.

### Data post processing and statistical analyses

#### CYTOKINE DATA

Principal component analysis was performed using *prcomp* in R *stats* package version 4.1.2 with the inoculated mice (*n*=12) and PBS mice (*n*=4) as observations and the log_10_ transformed cytokine abundance (mean-centered and scaled to unit variance) as variables. The relatedness of mice based on their cytokine profiles were assessed using hierarchical clustering analysis on the Euclidean distance between mice and cytokines with average linkage using R *pheatmap* package version 1.0.12.

#### VOLATILE DATA

The data post processing steps are depicted in Figure S5. Before statistical analyses, compounds eluting prior to 358 s (acetone retention time) and siloxanes (i.e., chromatographic artifacts) were removed from the peak table. Missing values were handled using RepHM imputation in the R *MetabImpute* package version 0.1.0, as follows: peaks that were present in only one of the three technical replicates were imputed to zero for that replicate, while missing values for peaks that were present in two out of three technical replicates were imputed to half of the minimum value within the replicates (as described in (61)). The relative abundances of compounds across chromatograms were normalized using probabilistic quotient normalization (PQN) (62) in R version 4.1.2. The data were log_10_ transformed, and geometric means of the technical triplicates were calculated. Principal component analyses were performed using *prcomp* in R *stats* package version 4.1.2 with the geometric means of the technical replicates as observations and the absolute peak intensities (mean-centered and scaled to unit variance) as variables. Kendall correlation analysis was performed correlating peak intensities to cytokine abundance. Analytes with Kendall correlation scores of –0.3 > τ > 0.3 and P < 0.05 were considered to be significantly correlated. The relatedness of samples based on their volatile metabolomes were assessed using hierarchical clustering analysis using R *pheatmap* package version 1.0.12.

## Data availability

Metabolomic data (chemical feature peak areas and retention time information) included in this study are available at the NIH Common Fund’s National Metabolomics Data Repository (NMDR) website, the Metabolomics Workbench, at www.metabolomicsworkbench.org, where it has been assigned project ID PR0001064 and study ID TBD (https://doi.org/TBD).

## ACKNOWLEDGEMENTS

Funding for this work was provided by the Arizona Biomedical Research Centre award ADHS18-198861 (H.D.B.).

We thank Bilal Ali for ascertaining the linear relationship between the 624Sil column and nonpolar stationary phase, which lead to the calculated 624Sil equation used for identifying level 2 compounds in this study.

