## Supplemental Figures and Tables for "Volatile metabolites in lavage fluid are correlated to Valley fever disease severity in murine model lung infections"

Running Title: Volatiles correlate to Valley fever infections in mice

[*] Present address: Marley C. Caballero Van Dyke, Department of Microbiology, University of Texas Southwestern Medical Center, Dallas, Texas, USA

Heather L. Mead, The Translational Genomics Research Institute (TGen), Phoenix and Flagstaff, Arizona, USA

**Results**

**Table S1:** Table of fungal dissemination to mouse spleen and brain (CFU/ml) and cytokine abundance (pg/ml).

| | Dissemination (CFU/ml) | | Anti-Inflammatory (pg/ml) | | | | | Pro-inflammatory (pg/ml) | | | | | | | | | | Multi-faceted (pg/ml) | | Chemokine (pg/ml) | | | | | | | | |
|---|---|---|---|---|---|---|---|---|---|---|---|---|---|---|---|---|---|---|---|---|---|---|---|---|---|---|---|---|
| **Mouse** | **Spleen** | **Brain** | **IL-4** | **IL-5** | **IL-10** | **IL-13** | **IL-22** | **IL-1β** | **IL-2** | **IL-6** | **IL-9** | **IL-12p70** | **IL-17** | **IL-18** | **IL-23** | **TNF-α** | **IFN-γ** | **IL-27** | **Gm-CSF** | **Eotaxin** | **IP-10** | **KC** | **MIP1α** | **MIP1β** | **MIP2** | **MCP-1** | **MCP-3** | **RANTES** |
| **PBS 1** | 0 | 0 | 9 | 6 | 18 | 35 | 7 | 6 | 21 | 10 | 15 | 14 | 9 | 11 | 16 | 17 | 14 | 8 | 8 | 19 | 22 | 12 | 47 | 23 | 9 | 13 | 20 | 57 |
| **PBS 2** | 0 | 0 | 8 | 5 | 21 | 33 | 7 | 5 | 26 | 10 | 17 | 12 | 9 | 11 | 25 | 26 | 20 | 7 | 7 | 19 | 24 | 10 | 46 | 18 | 9 | 12 | 20 | 45 |
| **PBS 3** | 0 | 0 | 9 | 6 | 21 | 30 | 7 | 6 | 25 | 13 | 17 | 14 | 11 | 11 | 19 | 19 | 18 | 7 | 8 | 20 | 25 | 11 | 45 | 18 | 9 | 12 | 20 | 51 |
| **PBS 4** | 0 | 0 | 9 | 6 | 20 | 35 | 7 | 6 | 23 | 11 | 19 | 15 | 10 | 11 | 19 | 20 | 18 | 7 | 8 | 24 | 24 | 11 | 45 | 23 | 9 | 13 | 21 | 60 |
| **RS 1** | 10 | 8 | 95 | 64 | 157 | 108 | 44 | 49 | 47 | 6640 | 44 | 233 | 264 | 436 | 60 | 537 | 78 | 26 | 206 | 212 | 130 | 422 | 2944 | 1037 | 1892 | 110 | 1931 | 96 |
| **RS 2** | 160 | 1 | 80 | 34 | 74 | 63 | 20 | 21 | 29 | 2734 | 27 | 104 | 108 | 214 | 35 | 160 | 44 | 15 | 96 | 118 | 98 | 210 | 722 | 279 | 249 | 35 | 738 | 77 |
| **RS 3** | 16 | 9 | 76 | 47 | 105 | 72 | 25 | 27 | 37 | 3828 | 33 | 140 | 157 | 268 | 47 | 238 | 56 | 19 | 116 | 134 | 140 | 283 | 1595 | 600 | 438 | 44 | 1022 | 71 |
| **RS 4** | 0 | 0 | 29 | 13 | 31 | 50 | 10 | 10 | 26 | 836 | 21 | 39 | 44 | 67 | 25 | 57 | 30 | 10 | 26 | 39 | 46 | 108 | 560 | 167 | 61 | 18 | 225 | 56 |
| **RS 5** | 0 | 0 | 9 | 6 | 14 | 33 | 7 | 6 | 18 | 38 | 14 | 13 | 10 | 13 | 12 | 15 | 12 | 7 | 8 | 23 | 25 | 15 | 75 | 25 | 11 | 12 | 36 | 51 |
| **RS 6** | 0 | 0 | 11 | 6 | 18 | 30 | 7 | 6 | 20 | 16 | 16 | 14 | 10 | 12 | 14 | 16 | 18 | 7 | 8 | 22 | 70 | 13 | 55 | 22 | 11 | 13 | 126 | 47 |
| **Sil 1** | 31 | 7 | 128 | 64 | 41 | 57 | 17 | 16 | 27 | 1259 | 25 | 59 | 77 | 140 | 21 | 73 | 27 | 11 | 46 | 179 | 154 | 64 | 504 | 365 | 21 | 37 | 3101 | 63 |
| **Sil 2** | 31 | 0 | 259 | 88 | 34 | 67 | 16 | 16 | 27 | 1122 | 27 | 54 | 55 | 170 | 18 | 67 | 18 | 13 | 46 | 168 | 599 | 73 | 689 | 495 | 27 | 66 | 6295 | 80 |
| **Sil 3** | 620 | 3 | 213 | 112 | 42 | 62 | 23 | 17 | 27 | 1309 | 27 | 66 | 55 | 171 | 23 | 114 | 42 | 14 | 49 | 178 | 376 | 73 | 950 | 540 | 44 | 72 | 5602 | 69 |
| **Sil 4** | 13 | 0 | 72 | 20 | 32 | 48 | 11 | 12 | 27 | 1045 | 20 | 48 | 51 | 100 | 21 | 65 | 24 | 10 | 35 | 107 | 93 | 113 | 600 | 280 | 23 | 23 | 1553 | 59 |
| **Sil 5** | 30 | 7 | 28 | 10 | 20 | 33 | 8 | 7 | 22 | 238 | 18 | 25 | 20 | 34 | 21 | 32 | 22 | 8 | 14 | 51 | 50 | 37 | 190 | 87 | 13 | 15 | 200 | 48 |
| **Sil 6** | 9400 | 8 | 329 | 333 | 83 | 91 | 44 | 33 | 43 | 2978 | 39 | 129 | 240 | 316 | 35 | 278 | 50 | 20 | 107 | 687 | 826 | 163 | 2489 | 1897 | 118 | 135 | 5758 | 92 |

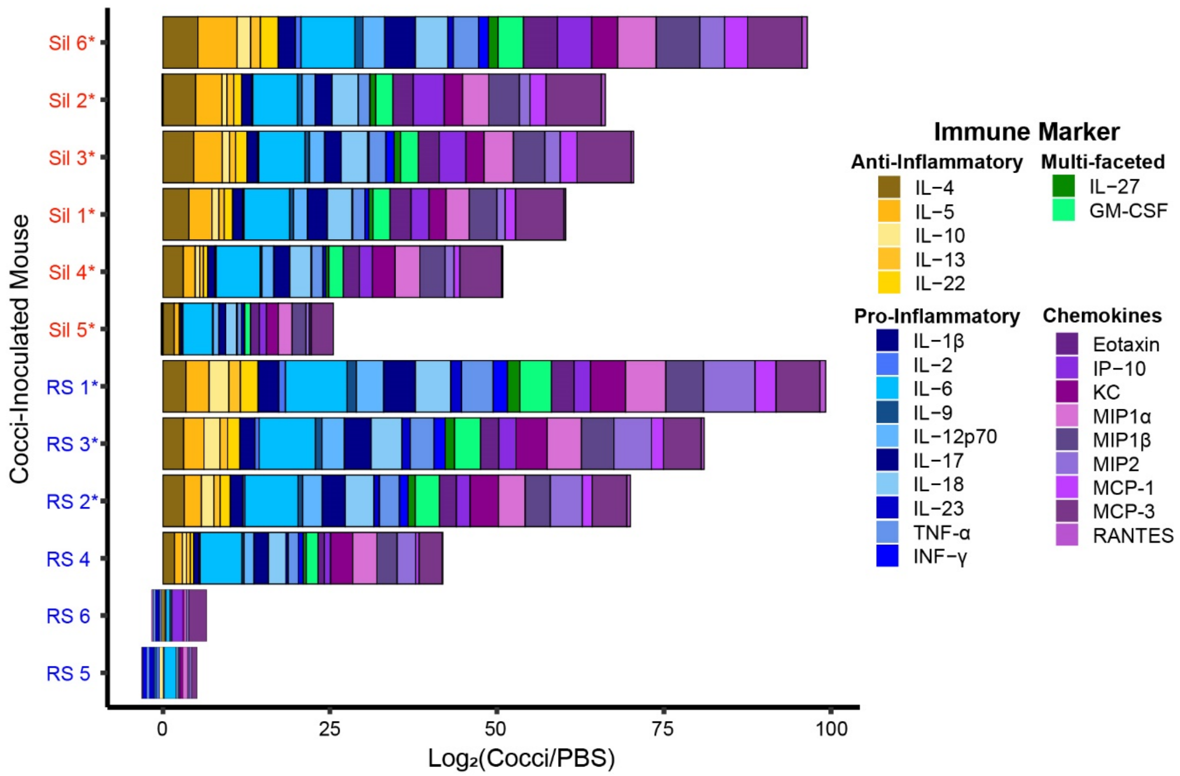


**Figure S1:** $Log_2$ fold change of cytokine abundances of Cocci-inoculated mice (*C. immitis* strain RS (blue) and *C. posadasii* strain Silveira (Sil; red)) relative to the mean cytokine abundances of the PBS-inoculated mice. Mice with disseminated disease are indicated with an asterisk (*). Immune markers are color-coded by type.

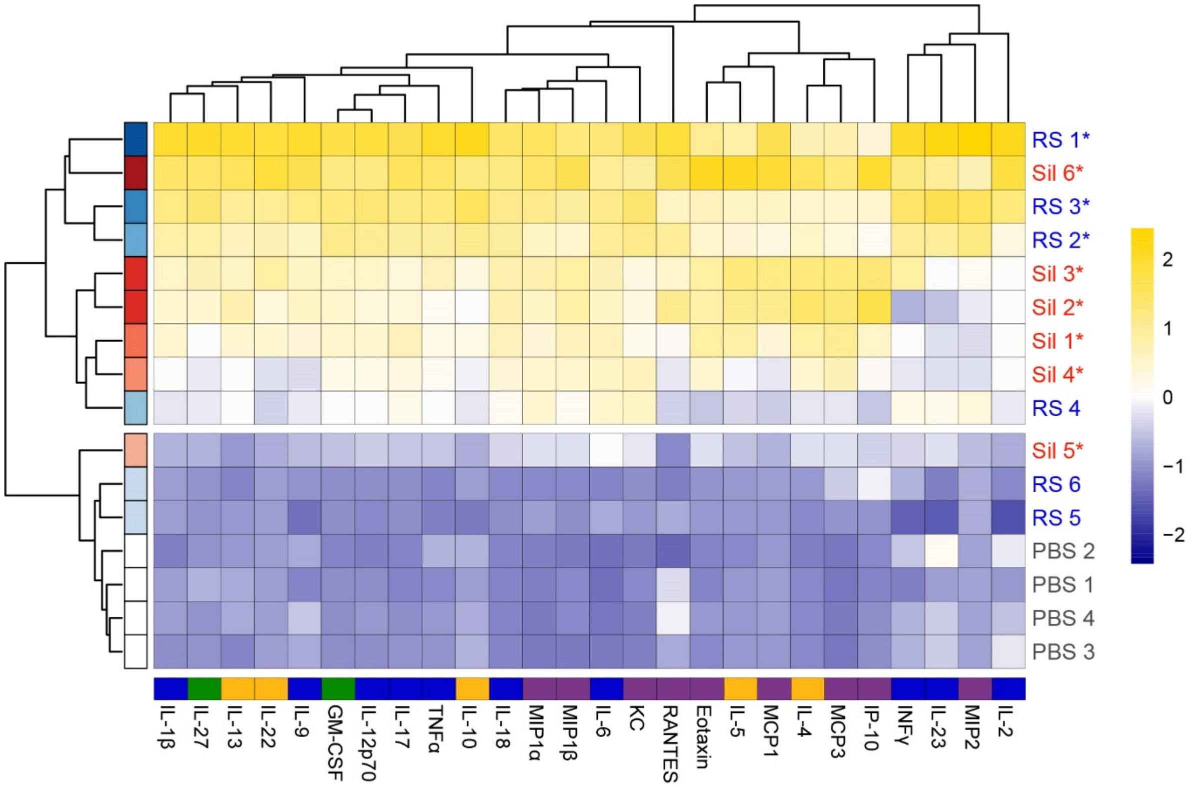


**Figure S2:** Hierarchical clustering analysis (HCA) of 12 Cocci-inoculated and 4 PBS-inoculated mice (rows) based on the relative abundances of 26 cytokines (columns). Clustering of mice and cytokines uses Euclidian distance with average linkage. Mice are color-coded by strain (blue = *C. immitis* RS; red = *C. posadasii* Sil) and a color gradient indicating total cytokine abundance, with darker color meaning higher abundance; disseminated disease is indicated with an asterisk (*). Cytokines are color-coded by type (anti-inflammatory in yellow, pro-inflammatory in blue, multi-faceted in green, and chemokines in purple), and their abundances (mean-centered and scaled to unit variance) are represented in the heat map

**Table S2:** See **Supplementary Excel File**. Table of 91 *Coccidioides* VOCs detected in the headspace of mouse bronchoalveolar lavage fluid samples.

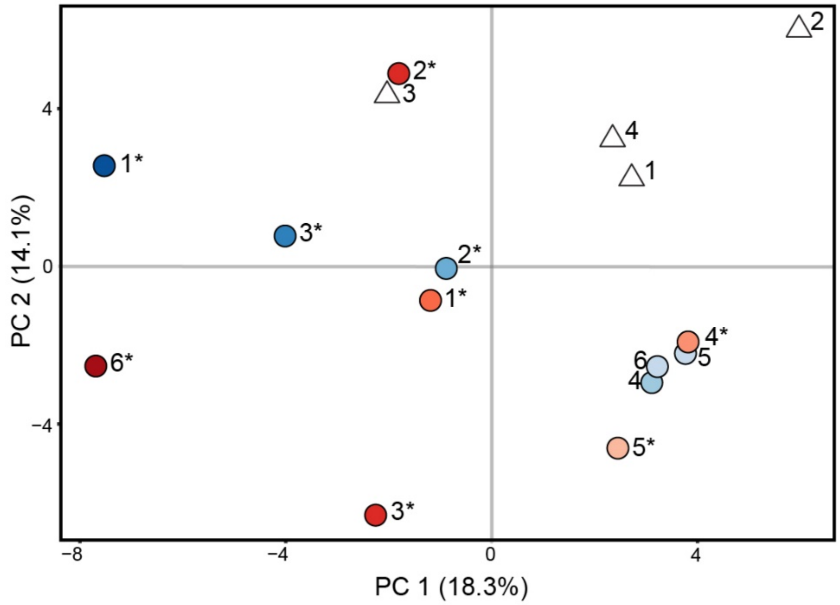


**Figure S3:** Principal component analysis (PCA) score plot using 91 volatile organic compounds (VOCs) detected in the BALF of mice inoculated with *C. immitis* RS (blue circles), *C. posadasii* Silveira (red circles) or PBS (white triangles) as features. The color gradient, darkest to lightest, indicates total BALF cytokine abundance, highest to lowest. Disseminated disease is indicated with an asterisk (*).

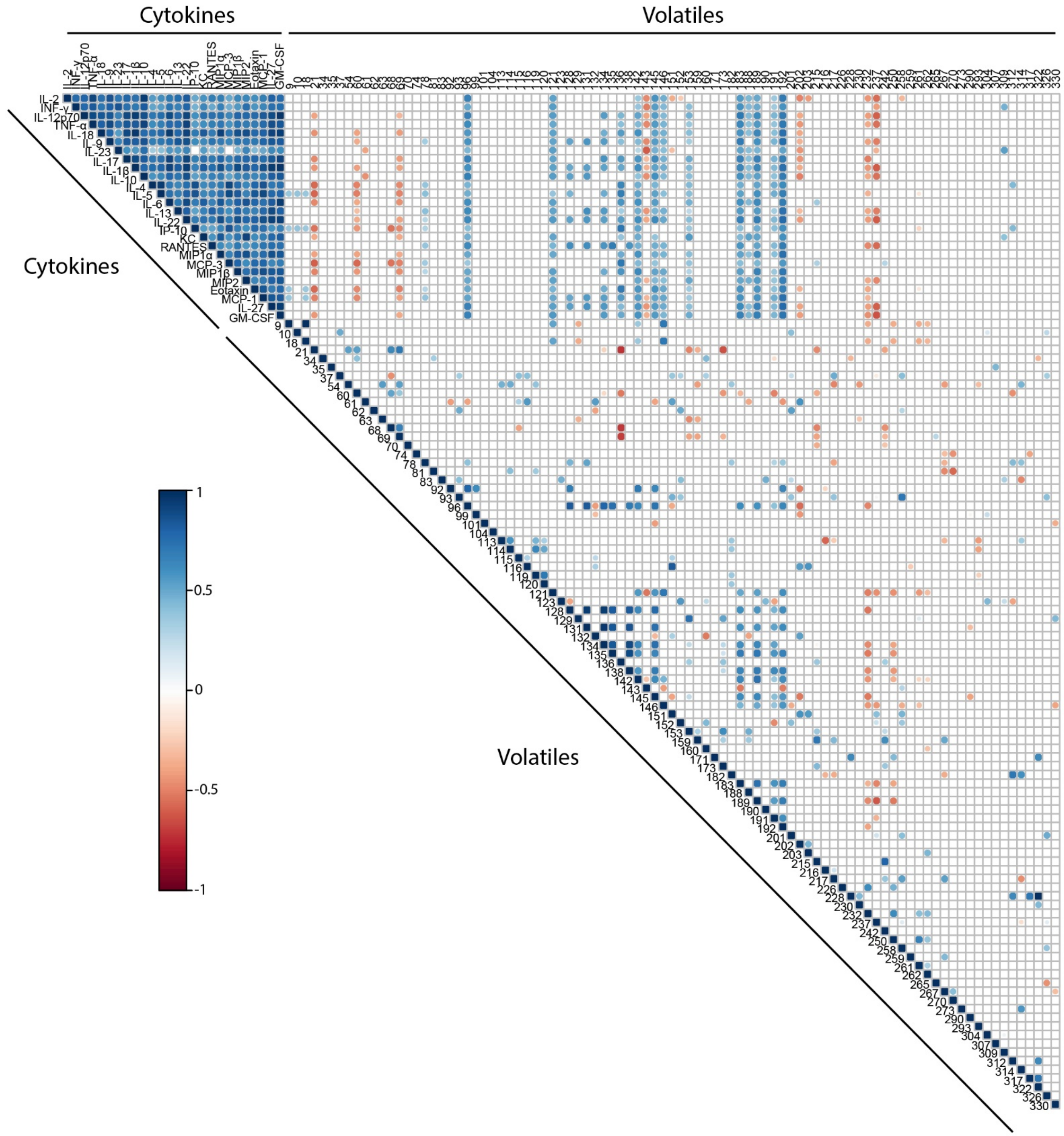


**Figure S4:** Kendall correlation plot of the abundance of 26 cytokines and 91 volatile organic compounds (VOCs) in 12 Cocci-inoculated and 4 PBS mice. Circles indicate statistically significant correlations ($P < 0.05$), with positive correlations in blue ($\tau > 0.3$), negative correlations in red ($\tau < -0.3$), and darker colors and larger sizes indicating a stronger correlation.

**Methods**

**Table S3:** Parameters for HS-SPME and GC×GC-TOFMS analysis, and data processing and alignment

| Autosampler Method | |
|---|---|
| Instrument description | Gerstel® MPS Pro® |
| Software description | Gerstel® Maestro® (version 1.5.3.2) |
| **Sampling Parameters** | |
| Cooled tray temperature | 4 °C |
| Solid-phase microextraction (SPME) | Manufacturer: Supelco®<br>Fiber type: PDMS/CAR/DVB (2 cm; 50/30 µm) |
| Incubation time | 2 min |
| Agitator parameters, incubation | Temperature: 50 °C<br>On time: 10 s<br>Off time: 1 s<br>Speed: 600 rpm |
| Agitation, sampling | On |
| Vial penetration | 21 mm |
| Extraction time | 10 min |
| Injection penetration | 67 mm |
| Desorption time | 180 s |
| **Inlet (CIS) Parameters** | |
| Initial temperature | 250 °C |
| Equilibrium time | 0.05 min |
| Initial time | 0.10 min |
| Ramp rate | 12 °C·s$^{-1}$ |
| End temperature | 250 °C |
| Hold time | 10.5 min |

| GC×GC Method | |
|---|---|
| Instrument description | Agilent® 7890B |
| Column configuration | Column 1: Rxi®-624Sil MS, 60 m × 0.25 mm × 1.4 *µ*m<br>Column 2: Stabilwax®, 1 m × 0.25 mm × 0.5 *µ*m |
| Carrier gas | Helium, 2 mL·min$^{-1}$ (constant) |
| Front inlet type | Gerstel® |
| Front inlet mode | Splitless |
| Front inlet septum purge flow | 1 mL·min$^{-1}$ |
| Front inlet septum purge time | 300 s |
| Front inlet purge flow | 50 mL·min$^{-1}$ |
| Front inlet total purge flow | 52 mL·min$^{-1}$ |
| Oven equilibration time | 5 s |
| Primary oven temperature ramp | Initial temperature: 35 °C<br>Initial time: 0.5 min<br>Ramp rate: 5 C·min$^{-1}$ |

| | |
|---|---|
| | Final temperature: 230 °C<br>Hold time: 5 min |
| Secondary oven temperature offset | +5 °C (relative to primary oven) |
| Modulator temperature offset | +15 °C (relative to secondary oven) |
| Modulation timing | Modulation period: 2.00 s<br>Hot pulse time: 0.50 s<br>Cold pulse time: 0.50 s |
| Transfer line temperature | 250 °C |
| **Mass Spectrometry Method** | |
| Instrument description | LECO® Pegasus® 4D |
| Use GC method total time for MS method total time | Yes |
| Acquisition delay | 180 s |
| Filament active time | 180 s to end of run |
| Start mass/End mass | 35/400 |
| Acquisition rate | 100 spectra·$s^{-1}$ |
| Optimized voltage offset | +50 V |
| Electron energy | -70 eV |
| Ion source temperature | 250 °C |
| **Data Processing Method** | |
| Software description | LECO® ChromaTOF® and Statistical Compare (version 4.71.0.0) |
| Baseline tracking/Offset | Entire run/0.5 (through middle of noise) |
| Data points averaged for smoothing | Auto |
| First dimension peak width | 12 slices |
| Mass spectral match required to combine peaks | 600 |
| Second dimension peak width | 0.15 |
| Min. subpeak signal-to-noise (S/N) for retention | 6 |
| Integration approach | Traditional |
| Peak finding | S/N: 50<br>Number of apexing masses: 2 |
| Mass spec libraries for searching | NIST® 2014 |
| Mass to use for area/height calculation | Unique mass |
| Alignment analyte match criteria | Spectral match mass threshold: 10<br>Minimum spectral similarity match: 600<br>Max. number of modulation periods apart: 1<br>Max. retention time difference (s): 0.1<br>S/N for second peak find: 5 |
| Criteria for inclusion of analytes | Min. number of samples that contain analyte: 1 |

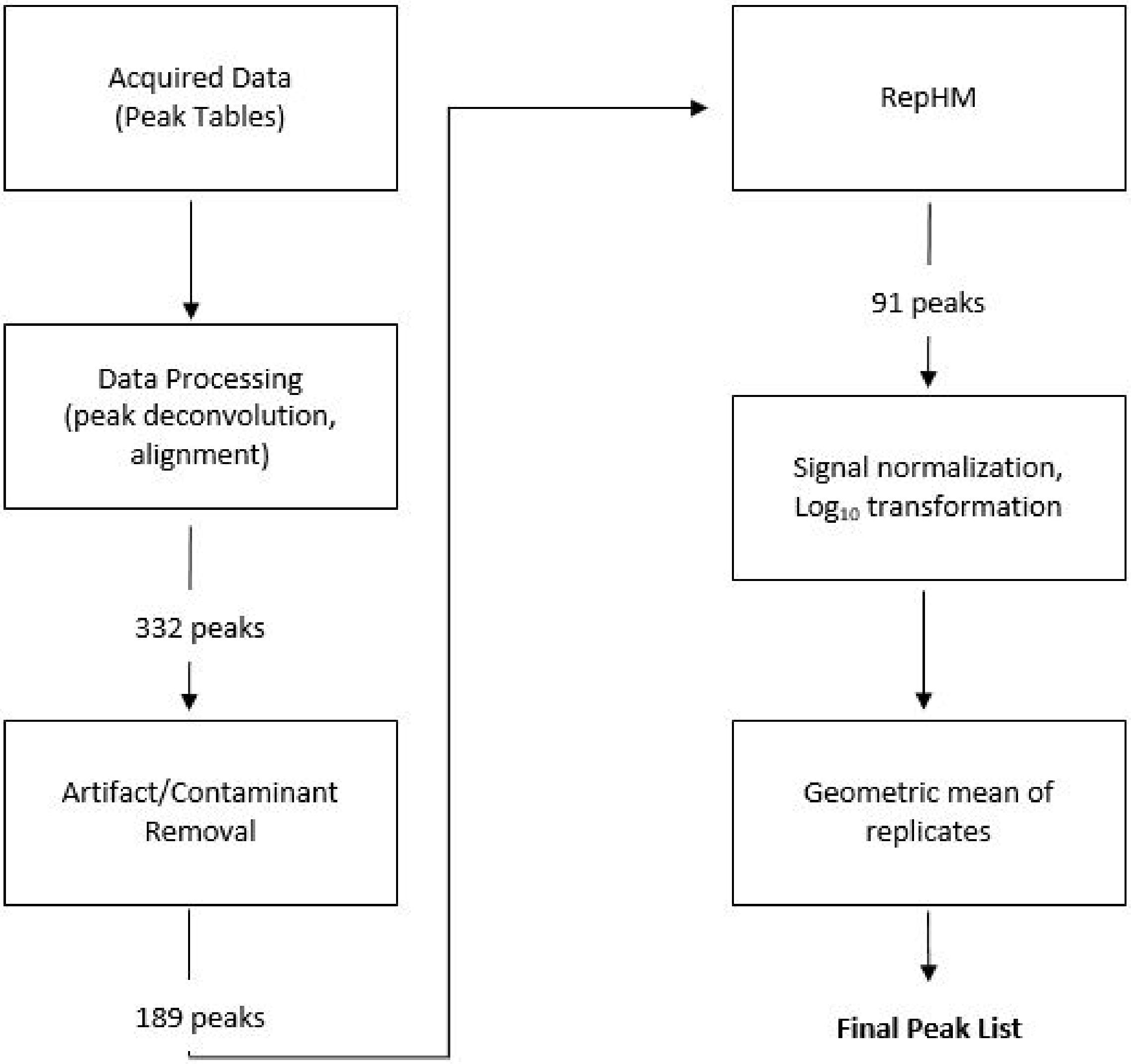


**Figure S5:** Data post processing workflow
